# Comparative transcriptomics reveal that adaptive evolution in immune genes drives the local adaptation and speciation of schizothoracine fish

**DOI:** 10.1101/714881

**Authors:** Delin Qi, Rongrong Wu, Yan Chao, Mingzhe Xia, Qichang Chen, Zhiqin Zheng

## Abstract

Transcriptomic information can increase our understanding of the molecular processes underlying speciation. The schizothoracine fish, the largest and most diverse taxon within the Qinghai-Tibetan Plateau (QTP) ichthyofauna, are widespread in drainages throughout the QTP. These fish thus serve as an ideal model group with which to investigate how molecular evolution drives local adaptation during speciation. Here, we performed an interspecific comparative analysis of the transcriptomes of 13 schizothoracine fish species, and identified the key positively selected genes (PSGs) associated with significantly enriched functions and metabolite pathway acting on the specific lineages (or species) in the schizothoracine fish. We generated 64,637,602–83,968,472 sequence reads per schizothoracine fish species using Illumina sequencing, yielding 95,251–145,805 unigenes per species. We identified 52 out of 2,064 orthologous genes as candidate genes, which have probably been subject to positive selection along the whole schizothoracine fish lineage. Nine of these candidate genes were significantly enriched in key GO functions and metabolite pathways, all of which were associated with the immune system. The lineage-specific evolution test showed species-specific differences among the nine candidate PSGs, probably due to ecological differences among drainages, as well as among micro-habitats in the same drainage (e.g., benthic and pelagic). Here, we provide evidence that the adaptive evolution of immune genes, along with the uplift of the QTP, allowed new schizothoracine species to colonize ecologically novel environments or to exploit vacant ecological niches during speciation.

Supplemental material available at FigShare: https://doi.org/10.25387/.

---

The processes and mechanisms of speciation have long been a major question in evolutionary biology (Funk et al. 2002; Rocha et al. 2005; Baack and Stanton 2005; Tonnis et al. 2005; Nosil et al. 2009; Malinsky et al. 2015). Local adaptations, driven by small-scale ecological niches, often result in adaptive phenotypes and the genetic divergence of geographically isolated populations (Baack and Stanton 2005; Nosil et al. 2009; Collin and Fumagalli 2011). The accumulation of genetic differences may lead to the formation of new taxon (Zhang et al. 2013; Collin and Fumagalli 2011; Nosil et al. 2009; Kirkpatrick and Barton 2006). Genetic divergences associated with local adaptations allow species to colonize ecologically novel environments or to exploit vacant ecological niches, and thus play an important role in speciation (Zhou et al. 2012; Yang et al. 2014; Kang et al. 2017; Tong et al. 2017; Cai et al. 2013; Qu et al. 2013). Freshwater fish, especially fish lineages distributed in different ecological drainage systems within a given region, have provided key insights into the mechanisms whereby molecular evolution drives local adaptation during speciation (Yang et al. 2016; Kang et al. 2017; Tong et al. 2017; Tong et al. 2015; Ma et al. 2015; Guan et al. 2014; Xu et al. 2017).

Schizothoracine fish (Teleostei: Cyprinidae) are the largest and most diverse taxon within the Qinghai-Tibetan Plateau (QTP) ichthyofauna, including more than 70 recognized species (Wu and Wu 1992; Chen and Cao 2000). Schizothoracine fish are distributed throughout the majority of the drainage basins in the QTP, 1500–5500 m above sea level (Qi et al. 2006; Wu and Wu 1992; Chen and Cao 2000). The distribution range of these fishes include the Yellow River, Yangtze River, Indus River, Mekong River, Tsangpo River, Salween River, Brahmaputra River, Qiadam Basin, and isolated lakes (Wu and Wu 1992; Chen and Cao 2000). These water bodies represent rich ecological diversity and complexity (Wu and Wu 1992; Qi et al. 2012). From an evolutionary perspective, the schizothoracine fish can be divided into three groups: primitive, specialized, and highly specialized, corresponding with three particular stages of QTP geological evolution (Cao et al. 1981). It has been demonstrated that the uplifting of the QTP from 50 MYA had a profound effect on paleo-drainage and stimulated many paleo-environmental changes in the plateau, thus promoting the speciation of the schizothoracine fish endemic to this region and shaping current species distributions (Qi et al. 2012; Li et al. 2013). Due to the extensive ecological diversity of the distribution drainages, and because speciation events occurred during the short period of QTP uplift, the schizothoracine fish is an ideal model group with which to investigate how molecular evolution drives local adaptation during speciation. Previous studies, using transcriptomic comparisons between fish species endemic to the QTP and other fish species endemic to lowland areas, have provided evidence of the genetic adaptation of schizothoracine fish to high-altitude environments (Yang et al. 2014; Tong et al. 2015; Tong et al. 2017). However, previous studies have only focused on the adaptive genetic differences between QTP schizothoracine fish and low-altitude fish species; the potentially adaptive genetic differences among the schizothoracine fish remain unclear. In addition, previous studies have only included a single schizothoracine species, which may have been insufficient to fully characterize the genetic signals of local adaptation during speciation in this group.

In this study, we sequenced, assembled, and merged the transcriptomes of four tissues (head kidney, muscle, brain, and liver) from 13 schizothoracine species. We then conducted a comparative transcriptomic analysis to identify key positively selected genes (PSGs) associated with significantly enriched functions and metabolite pathways acting on specific lineages (or species) of schizothoracine fish. We aimed to provide evidence of the genetic adaptation of schizothoracine fish to their local aquatic environment, leading to speciation.

## MATERIALS AND METHODS

### Experimental Animals

Fish were collected from across their ranges using gill nets or cast nets (Figure 1). During field sampling, a blow to the head was used to stun fish. The liver, muscle, brain, and head kidney were then removed and frozen in liquid nitrogen until use. The specimens used in this study included 13 species, representing the primitive, specialized, and highly specialized groups of schizothoracine fishes. All of the procedures involving animals followed the guidelines of, and were conducted under the approval of, the Animal Care and Use Committee, Qinghai University, China.

**Figure 1.**
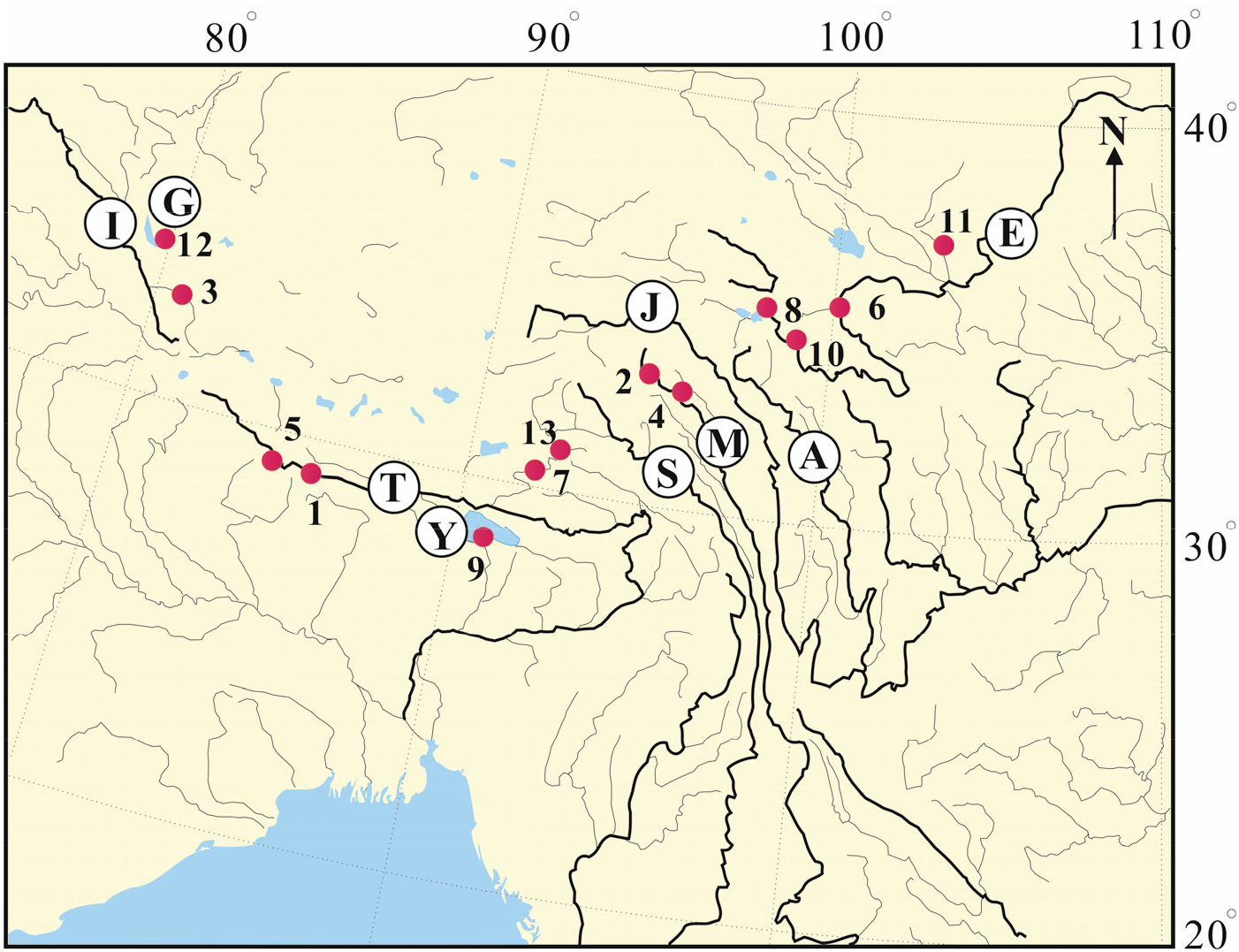
Location of fish sampling sites. Sites: A, Yalung River; E, Yellow River; G, Bangongco Lake; I, Indus River; J, Jinsha River; M, Mekong River; S, Salween River; T, Tsangpo River; Y, Yomzho Lake. Species sampled: 1, *Schizothorax macropogon*; 2, *Schizothorax lissolabiatus*; 3, *Schizothorax labiatus*; 4, *Schizothorax lantsangensis*; 5, *Ptychobarbus dipogon*; 6, *Gymnodiptychus pachycheilus*; 7, *Oxygymnocypris stewartii*; 8, *Gymnocypris eckloni*; 9, *Gymnocypris waddelli*; 10, *Platypharodon extremus*; 11, *Schizopygopsis pylzovi*; 12, *Schizopygopsis stoliczkai*; 13, *Schizopygopsis younghusbandi*.

### RNA extraction, Illumina library preparation, and sequencing

Total RNA was isolated from each sample using the Ambion Magmax-96 total RNA isolation kit (Life Sciences, USA), following the manufacturer’s instructions. RNA degradation and contamination was monitored using 1% agarose gels. RNA purity was checked using a Nano Photometer spectrophotometer (IMPLEN, CA, USA). RNA concentration was measured using Qubit RNA Assay Kits in a Qubit 2.0 Flurometer (Life Technologies, CA, USA). A single pooled RNA sample, containing equal volumes of RNA from each organ (liver, muscle, brain and head kidney), was prepared for each species. We used 3.0 μg of each pooled RNA sample used to prepare one Illumina sequencing library per species.

Sequencing libraries were generated using the NEBNext Ultra RNA Library Prep Kit for Illumina (NEB, USA), following manufacturer’s instructions. Index codes were added to attribute sequences to each sample. PCR products were purified using an AMPure XP system (Beckman Coulter, USA) and library quality was assessed with a Bioanalyzer 2100 (Agilent). The index-coded samples were clustered with a cBot Cluster Generation System using a TruSeq PE Cluster Kit v3-cBot-HS (Illumina), following the manufacturer’s instructions. After cluster generation, the library preparations were sequenced on an Illumina Hiseq 2500 platform, generating 125 bp paired-end reads. All sequence reads were deposited in the National Center for Biotechnology Information (NCBI) Sequence Read Archive database under bioproject number PRJNA494936.

### Quality control and de novo assembly

Raw data (raw reads) were sorted by individual species and processed using self-written Perl scripts. In this step, data (reads) were cleaned by removing reads containing adapter sequences, reads containing ploy-N sequences, and low-quality reads. All of the downstream analyses were based on high-quality clean data. De novo transcriptome assembly was performed with the short read assembly program Trinity (Grabherr et al. 2011), with min_kmer_cov set to 2 by default and all of the other parameters set to default (Grabherr et al. 2011). In brief, reads of a certain length with overlapping areas were first joined to form longer fragments (contigs) without gaps. Then, paired-end reads were mapped back to contigs. Finally, the contigs were connected until sequences could no longer be extended. These sequences were termed unigenes.

### Transcriptome annotation

All of the unigenes from the 13 schizothoracine fish were annotated against the NCBI non-redundant (Nr) protein database (ftp://ftp.ncbi.nih.gov/blast/db/) with BLAST (Altschul et al. 1997), setting an E-value cut-off of 1E-5. We also used BLAST to align and annotate unigene sequences against the nucleotide (Nt) database (ftp://ftp.ncbi.nih.gov/blast/db/), the Clusters of Orthologous Groups of proteins (KOG) database (ftp://ftp.ncbi.nih.gov/pub/COG/KOG/), Swiss-Prot (http://www.uniprot.org/), the Kyoto Encyclopedia of Genes and Genomes (KEGG) database (http://www.genome.jp/kegg/), and the Gene Ontology (GO) database (http://www.geneontology.org/). The results of the BLAST search against the GO database were imported into Blast2GO 3.2 (Gotz et al. 2008) for GO term mapping. The clusters of orthologous groups (COG) database were then used to identify putative functions for the unigenes, based on known orthologous gene products (Tatusov et al. 2003). The KEGG pathways were analyzed using the online KEGG Automatic Annotation Sever (http://www.genome.jp/kegg/kaas/), with the bi-directional best-hit method and an E-value cutoff of 1E-10.

### Ortholog identification and sequence alignment

Translated amino acid sequences from the 13 schizothoracine fish were used to construct a database, along with sequences from another two fish species: zebrafish (*Danio rerio*) and medaka (*Oryzias latipes*), obtained from the Ensembl database (release 89). Next, we used performed a self-to-self BLASTP against all of the amino acid sequences, with a E-value cutoff of 1e^−5^; hits with identity < 30% and coverage < 30% were removed. One-to-one orthologs between 13 schizothoracine fish were determined using OrthoMCL v2.0.9 software (Li et al. 2003) with default settings. Finally, putative single copy orthologs across the 13 schizothoracine fish were obtained. For genes with multiple transcripts, the longest transcript was chosen. Each orthologous gene set was aligned using MUSCLE v. 3.8.31 (Edgar 2004), and trimmed using Gblocks (Castresana 2000) with the parameter “–t = c”. We deleted all of the gaps and ambiguous bases (“N”) from the alignments to reduce the effects of these elements on positive selection inference. After this deletion process, trimmed alignments shorter than 150 bp (50 codons) were discarded.

### Phylogeny construction and PSGs identification across the schizothoracine fish lineage

We constructed a phylogenetic tree of the 13 schizothoracine fish using the concatenated orthologous sequences using a maximum-likelihood (ML) analysis in MEGA 6.0 (Tamura et al. 2013). The ML tree was then used in a PAML analysis (Yang 2007). We examined the schizothoracine fish phylogeny for evidence of natural selection by comparing non-synonymous/synonymous substitution ratios (ω = Ka/Ks), where ω = 1, <1, and>1 indicated neutral evolution, purifying selection, and positive selection, respectively (Yang and Nielsen 2002; Yang et al. 2005). To determine how many orthologous genes had undergone positive selection across the whole lineage of schizothoracine fishes, we applied site-specific ML models using the codeml in PAML (version 4.7) (Yang 2007). Site-specific models that allowed ω to vary among sites were used to detect site-dependent evolution across the schizothoracine fish lineage. Thus, all the putative single-copy genes identified in the 13 schizothoracine fishes were separately tested for positive selection using the neutral model (M7) and the selection model (M8).

### PSG GO enrichment and KEGG pathways analysis

GO functional enrichment and KEGG pathway analyses were performed for genes shown to be under positive selection across the lineage of the schizothoracine fishes. GO enrichment analysis was performed using GOseq (Young et al. 2010), which is based on a Wallenius non-central hyper-geometric distribution. KEGG pathways were analyzed using KOBAS v2.0.12 (Xie et al. 2011), with FDR set to BH.

### Lineage-specific evolution of the key PSGs associated significantly enriched functions and metabolite pathways

Positive selection pressure on genes is one of the fundamental processes underlying adaptive changes in genes and genomes, resulting in evolutionary innovations and species differences (Marra et al. 2017). During speciation and dispersal, the different lineages (or species) of the schizothoracine fish have been subject to a variety of different selection pressures, including low temperatures, hypoxic conditions, intense ultraviolet radiation, and unique pathogens (Wu and Wu 1992; Qi et al. 2012). To investigate which of the key PSGs were associated with significantly enriched functions and metabolite pathways acting on specific lineages (or species), which may have driven the adaption of specific taxa to local environmental conditions, we used branch-specific ML models (Zhang 2005). Branch-specific models allow the ω ratio to vary among branches in a phylogeny, and are useful for detecting positive selection pressures acting on particular lineages. Thus, for the positively selected genes significantly enriched in GO terms and KEGG pathways, we evaluated the ω ratio for every lineage of the schizothoracine fish, using branch-specific ML models with branch labels.

## RESULTS

### Illumina sequencing and de novo transcriptome assembly

We generated 64,637,602–83,968,472 sequence reads for each of the 13 schizothoracine fish with Illumina sequencing. After trimming and filtering low-quality sequences, we obtained 62,487,860–81,238,722 high-quality clean sequence reads for each of the 13 species. Due to the absence of genome information for the schizothoracine fishes, we de novo assembled the sequence reads using Trinity, obtaining 129,982–207,887 for each of the 13 species. Each of these transcripts included 95,251–145,805 unigenes (Table 1). Detailed unigene information is summarized in Supplementary Material 1.

**Table 1.**
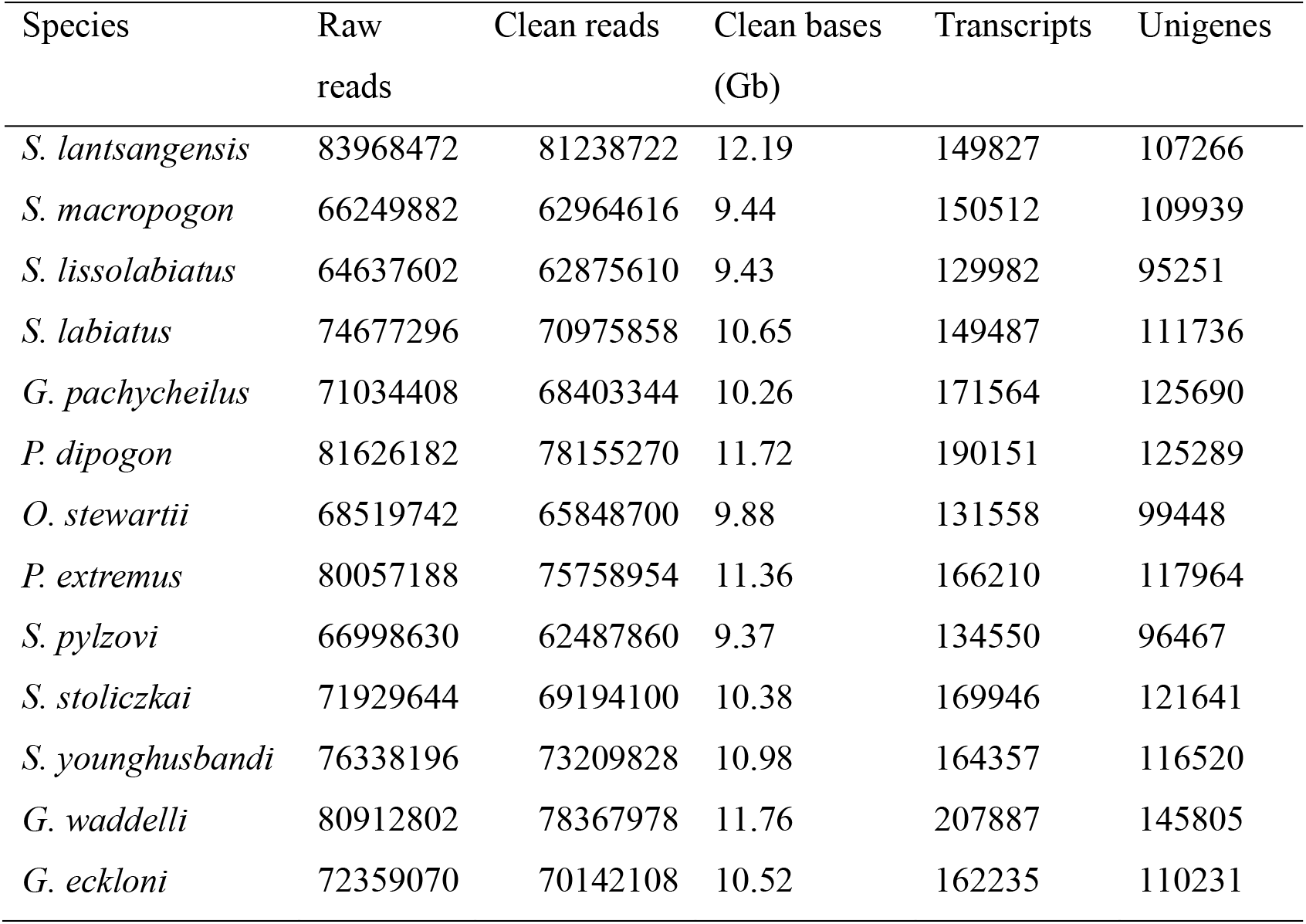
Summary of transcriptome data for 13 schizothoracine fish

### Unigene annotation

All of the unigene sets across the 13 schizothoracine fish were annotated based on similarity to sequences in six public databases. We found the largest number of unigene matches in the Nt database, ranging from 115,232 (79.03%: *Gymnocypris waddelli*) to 85,026 (89.26%: *Schizothorax lissolabiatus*). We found that 81.43%–90.94% of the unigenes from each species had a match in at least one database. We used the unigenes BLAST hits from the Nr database for subsequent analyses, as the Nr database had the maximum number of protein reference sequences per species (Table 2).

**Table 2.**
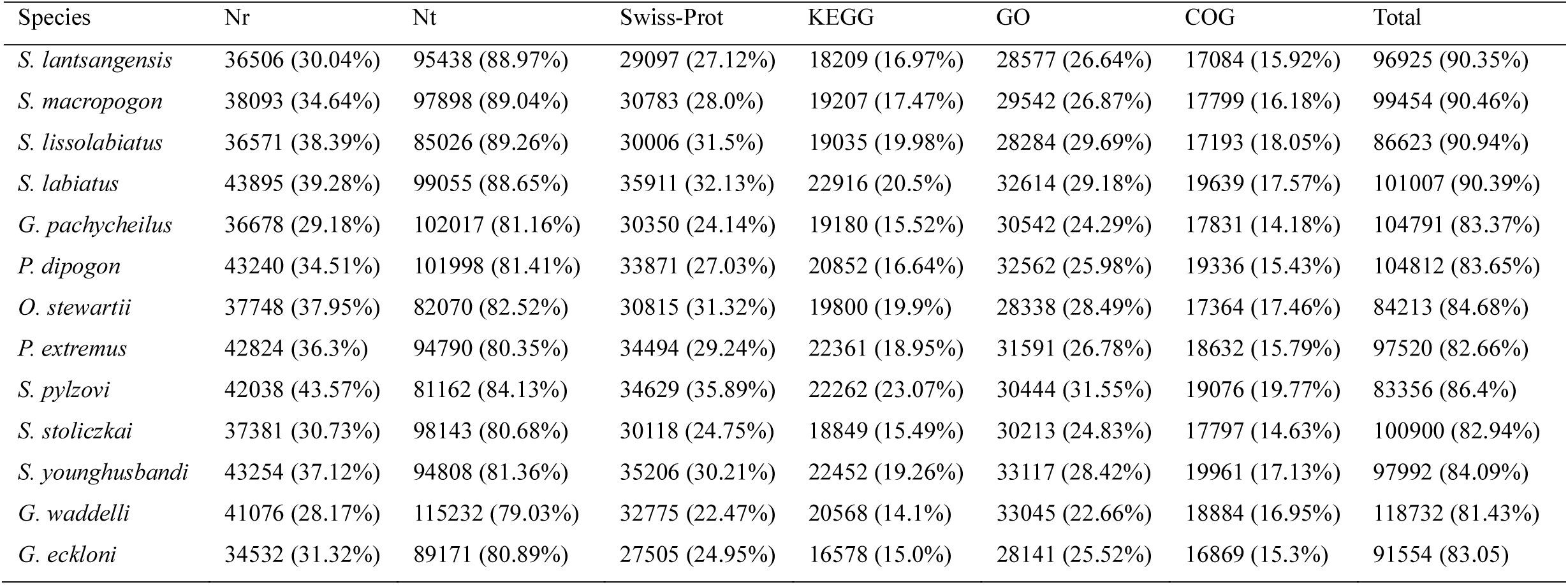
Functional annotation of unigenes for 13 schizothoracine fish.

### Functional annotation and classification

Based on sequence homology against the Nr database, the 28,141 (*Gymnocypris eckloni*) to 33,117 (*Schizopygopsis younghusbandi*) unigenes from the 13 schizothoracine fish were annotated to 56 GO categories (Table S2). Of these, there were 24 biological process (BP) sub-categories, 18 cellular component (CC) sub-categories, and 14 molecular function (MF) sub-categories. Binding (GO:0005488) was the most represented MF category; cellular process (GO:0009987) was the most represented BP category, and cell (GO:0005623) and cell part (GO:0044464) were the most represented CC categories (Supplementary Material 2).

We functionally classified 16,578 (*G. eckloni*) to 22,916 (*Schizothorax labiatus*) unigenes from the 13 schizothoracine fish into five KEGG categories and 32 functional sub-categories. Among the 32 subcategories, signal transduction were the most highly represented, followed by endocrine system, cell communication, nervous system, and immune system (Supplementary Material 3).

### Putative orthologs, phylogeny, and positively selected genes

We divided 477,274 proteins from the 13 schizothoracine fish and two other fish species (*D. rerio* and *O. latipes*) into 52,508 orthologous groups (gene families) using OrthoMCL (Li et al. 2003), following the self-self-comparison with BLASTP. After alignment and trimming for quality control, we compared the orthologous groups among the 13 fish species, and identified 2,064 putative single-copy genes in each fish species. These single-copy genes were used in the subsequent phylogenetic and evolutionary analyses. The ML phylogenetic tree, based on 2,064 orthologous genes, was well supported with 100% bootstrap values at all of the nodes (Figure 2). This ML tree was congruent with previous trees based on morphological characters (Wu and Wu 1992) and on mitochondrial genes (Saitoh et al. 2006; Qi et al. 2012).

**Figure 2.**
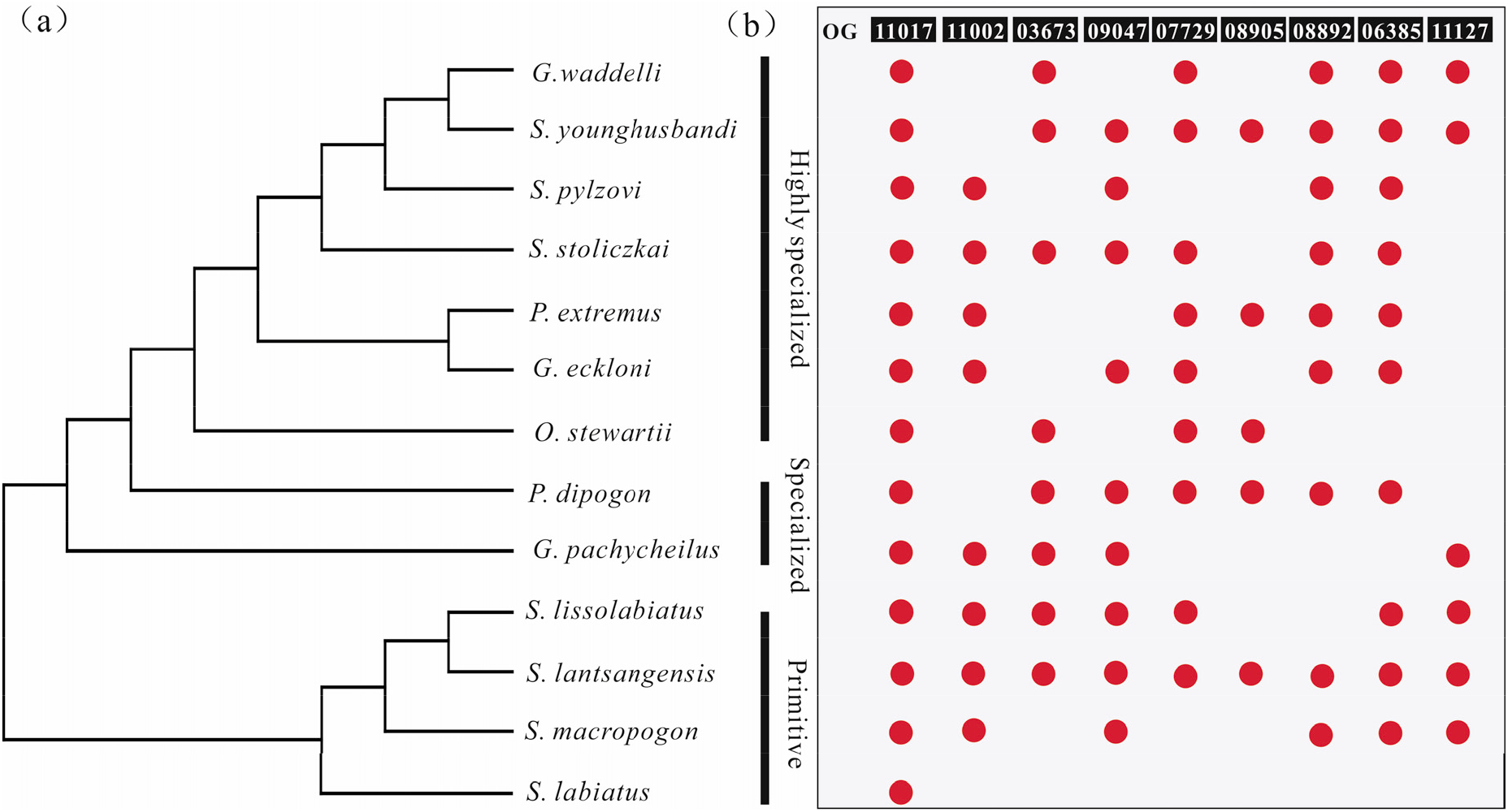
(a) Maximum likelihood (ML) phylogenetic tree, inferred based on 2064 orthologous genes. (b) Lineage-specific evolution test of the nine candidate positively selected genes (PSGs). Red dots represent genes with a Ka/Ks ratio >1 in a specific lineage.

Positive selection analysis pinpointed the genes that were associated with a functional or environmental shift. For the 2,064 orthologous genes that harbored both synonymous and non-synonymous substitutions, we identified 52 genes with Ka/Ks ratios >1, and 187 genes with Ka/Ks ratios between 0.5 and 1 (Table 3 and Figure 3). The 52 genes (with Ka/Ks ratios > 1) were considered candidate genes, that probably underwent positive selection along the schizothoracine fish lineage, and which may have influenced the adaptation of these fish to the aquatic environment of the QTP (Table 3 and Figure 3).

**Table 3.**
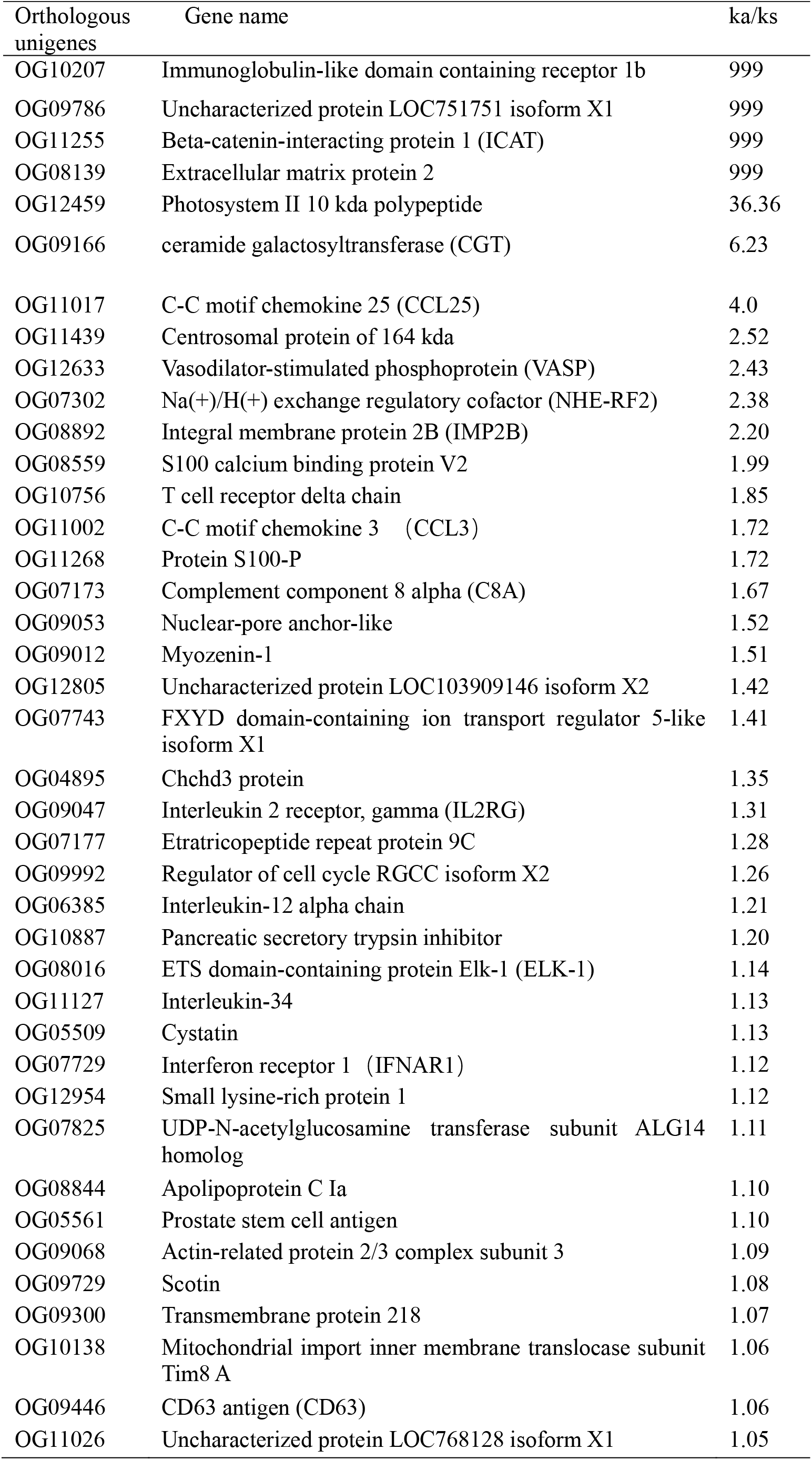
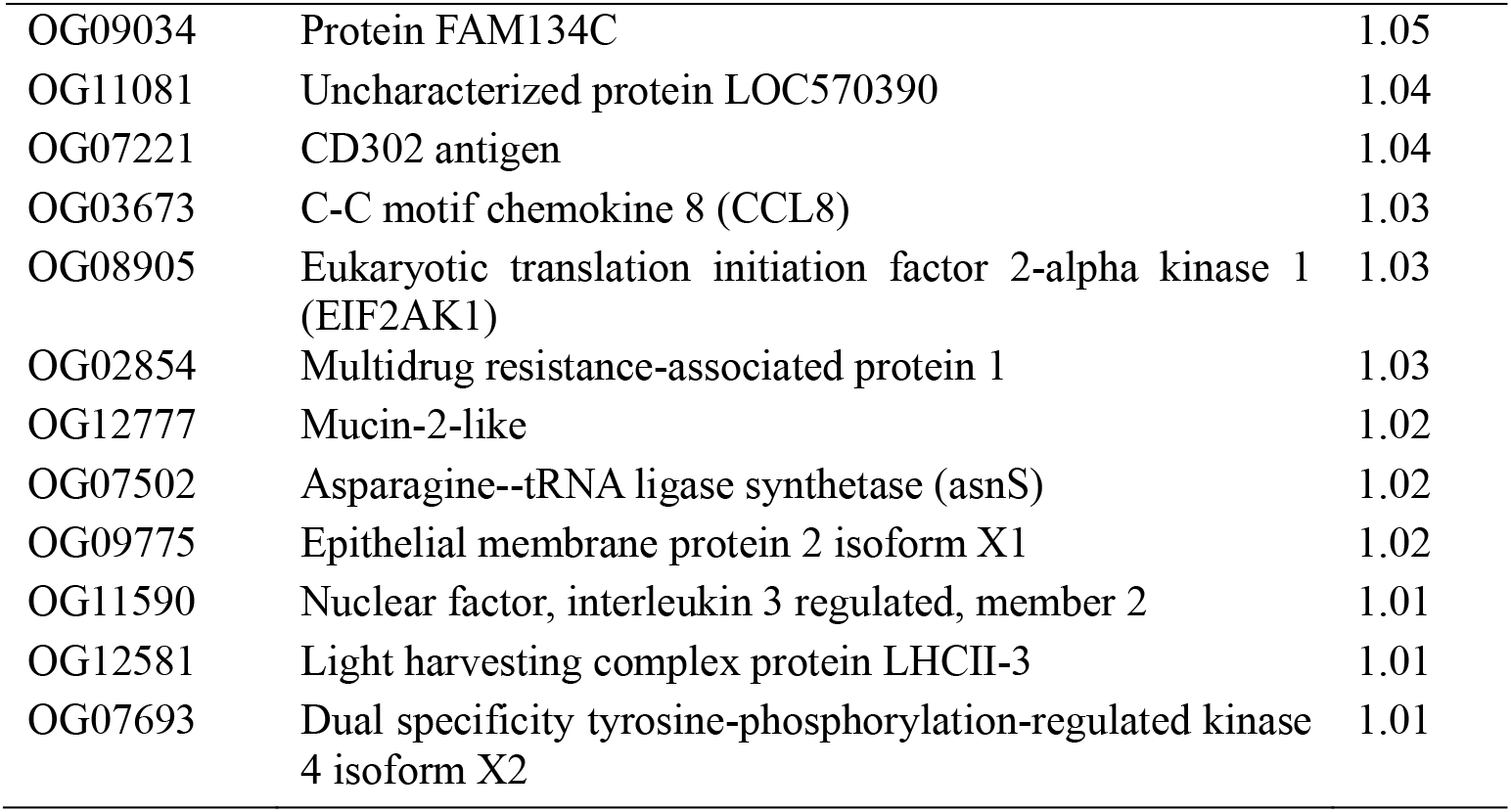
Identification of candidate genes under positive selection (Ka/Ks>1).

**Figure 3.**
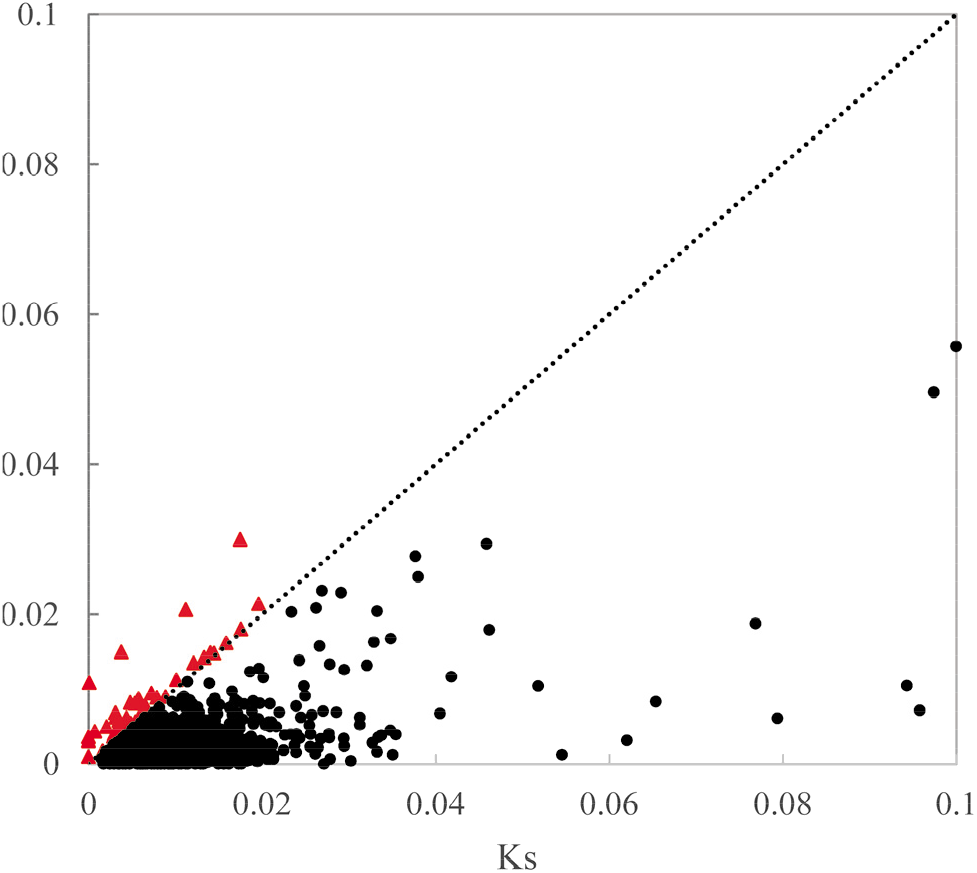
Plot of nonsynonymous (Ka) vs. synonymous (Ks) substitutions. The dotted line suggests neutrality, values above the line (red triangles) are subject to positive selection, and values below the line (black dots) are subject to purifying selection.

### PSG functions and metabolite pathways

To detect genes that might be involved in the adaptation to diverse aquatic environments in the QTP, we performed GO functional enrichment and KEGG pathways analysis on the 52 PSGs with orthologous unigenes.

Our GO enrichment analysis identified six PSGs (OG03673, OG08892, OG11017, OG06385, OG11002, and OG11127) that were significantly enriched in four molecular functions (cytokine activity, cytokine receptor binding, chemokine activity, and chemokine receptor binding), and one biological process (immune response; Table 4). KEGG enrichment analysis indicated that six PSGs (OG03673, OG09047, OG07729, OG11002, OG11017, and OG08905) were significantly enriched in two KEGG pathways (cytokine-cytokine receptor interaction and measles) (Table 5 and Figure 4). Thus, we identified nine candidate PSGs in the schizothoracine fish that may be involved in the adaptation to diverse aquatic environments. Notably, all nine of the candidate PSGs were related to fish innate and/or adaptive immunity.

**Table 4.** Key PSGs involved in the significantly enriched GO functions.

| GO accession | GO terms | GO type | Positively selected genes |
| --- | --- | --- | --- |
| GO:0005125 | Cytokine activity | MF | OG03673, OG08892, OG11017, OG06385, OG11002 |
| GO:0005126 | Cytokine receptor binding | MF | OG11017, OG11002, OG06385, OG11127, OG03673 |
| GO:0008009 | Chemokine activity | MF | OG11017, OG11002, OG03673 |
| GO:0042379 | Chemokine receptor binding | MF | OG03673, OG11017, OG11002 |
| GO:0006955 | Immune response | BP | OG03673, OG08892, OG11017, OG06385, OG11002 |

**Table 5.** Key PSGs involved in the significantly enriched KEGG pathways.

|  | KEGG pathway | ID | Positively selected genes |
| --- | --- | --- | --- |
| Immune system | Cytokine-cytokine receptor interaction | ko04060 | OG07729, OG11002, OG03673, OG09047, OG11017 |
| Immune system | Measles | ko05162 | OG07729, OG08905, OG09047 |
| Immune system | Chemokine signaling pathway | ko04062 | OG11002, OG03673, OG11017 |

**Figure 4.**
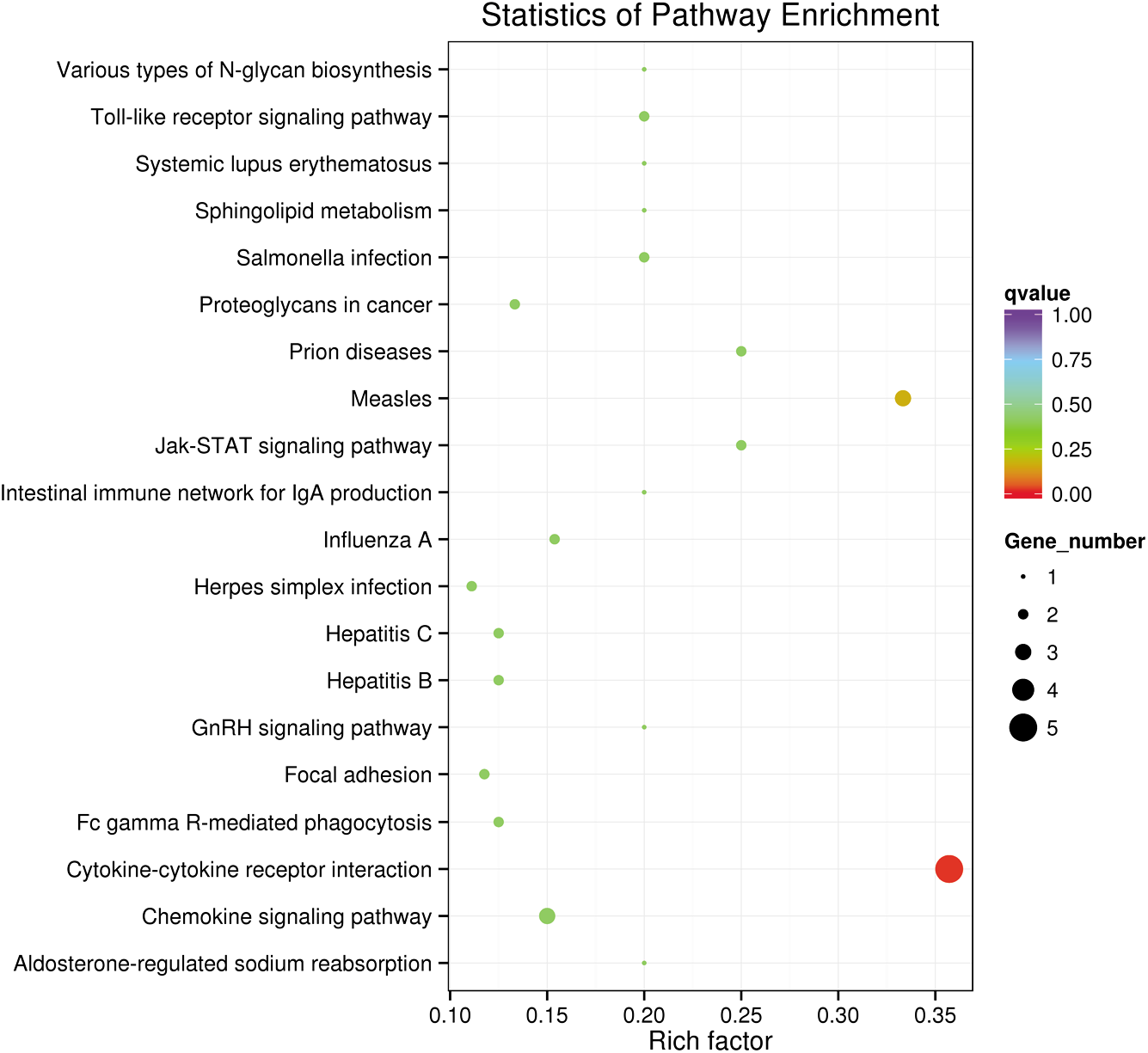
An overview of the KEGG pathways significantly enriched in the 52 PSGs. Specific pathways are plotted along the y-axis, and the x-axis indicates the enrichment factor. The size each colored dot indicates the number of candidate PSGs associated with each corresponding pathway: pathways with larger-sized dots contain a larger numbers of genes. The color of the each dot indicates the corrected *P*-value for the corresponding pathway.

The lineage-specific evolution test of the nine candidate PSGs indicated that different candidate PSGs acted more strongly on different specific lineages or species, as compared to other lineages or species (Figure 2). For example, of the 13 species, *S. lantsangensis* possessed the most PSGs (nine), followed by *Ptychobarbus dipogon*, *S. younghusbandi*, and *S. stoliczkai* (eight PSGs), while *S. labiatus* had the fewest PSGs (one).

## DISCUSSION

A better understanding of fish genetics will provide key insights into the adaptation of fish to diverse aquatic environments in the QTP, and thus increase our knowledge of the molecular mechanisms underlying the speciation, dispersal, and current distribution of the freshwater fish endemic to the QTP. Previous studies, comparing the transcriptomics of single QTP fish species of QTP to other low-land species, have identified a set of candidate genes related to energy metabolism and hypoxia response in freshwater fish in the QTP (Yang et al. 2014; Tong et al. 2017). Here, our aim was to characterize interspecific transcriptomic differences within the subfamily Schizothoracinae, and, more specifically, to identify the key PSGs involved in the significantly enriched functions and metabolite pathways acting on the specific lineages (or species). In this way, we aimed to investigate the genetic basis underlying the adaptation of this subfamily to local aquatic environments. As part of our study, we also sequenced and assembled merged-tissue transcriptomes of 13 schizothoracine species, providing an important genetic resource that was previously lacking.

Transcriptome quality is usually influenced by the method of library construction and the sequencing platform. Our assembly statistics indicated that the species with the greatest number of sequencing reads yielded more assembled transcripts, unigenes, and annotated open reading frames (ORFs). Our merged transcriptomes yielded 95,251–145,805 unigenes for each of the 13 schizothoracine species, which is consistent with a previous report of single-tissue RNA-seq data from *Gymnocypris przewalskii* (Zhang et al. 2015), but far greater than multiple-tissue merged transcriptomes from *Gymnodiptychus pachycheilus* (Yang et al. 2014), *G. przewalskii* (Tong et al. 2017), and glyptosternoid fish (Ma et al. 2015; Kang et al. 2017). The N50 values across the 13 species were similar, ranging between 1,030 and 1,567 kb (mean length ranging from 635–745 bp; Table S1). These N50 values were slightly smaller than those of *G.przewalskii* (Tong et al. 2017) but consistent with most recent comparative transcriptome studies in fish (Yang et al. 2014; Ma et al. 2015; Kang et al. 2017).

The number of Nr annotations was similar across the 13 species (ranging from 34,532 to 43,254; Table 2), and consistent with estimates for single-tissue RNA-seq data in *G. przewalskii* [17]. However, the number of Nr annotations recovered here was far greater than multiple-tissue merged transcriptomes produced for other fish species (Yang et al. 2014; Ma et al. 2015; Kang et al. 2017; Tong et al. 2017). Out of the 95,251–145,805 assembled unigenes from the 13 species, we annotated 81.43%–90.94% against six public databases. Unigenes without significant hits may be orphan genes, noncoding RNAs, untranslated transcripts, or misassembled transcripts. Thus, there might have been orphan genes specific to the schizothoracine fish (Yang et al. 2014), which might have rapidly evolved, taking indispensable biological roles and contributing to lineage-specific phenotypes and adaptations (Chen et al. 2013). Relatively few shared orthologs were identified because large interspecific comparisons are still somewhat nascent. Even so, we identified 2,064 orthologous genes in 13 schizothoracine species, which were fewer than that identified in two congeneric species of naked carp (Zhang et al. 2015), but the amount was consistent with similar comparative transcriptome studies in other fish species (Yang et al. 2014; Ma et al. 2015; Tong et al. 2017). Thus, we have here obtained a set of high quality transcriptomic resources, which are appropriate for further comparative genomic analyses when genome sequencing data are not available.

The schizothoracine fish (Teleostei: Cyprinidae) are the largest and most diverse taxon of the QTP ichthyofauna (Wu and Wu 1992; Chen and Cao 2000). Previous studies have demonstrated that the uplifting of the QTP from 50 MYA had a profound effect on drainage patterns and changed the environment of the plateau substantially, promoting the speciation of the schizothoracine fish endemic to the region (Li et al. 2013; Qi et al. 2015). Previous studies based on comprehensive transcriptomes have shown that genes associated with energy metabolism and the hypoxia response underwent significantly accelerated or adaptive evolution in the schizothoracine fish, as compared to other teleost fish from the plains (Yang et al. 2014; Tong et al. 2017). Here, we identified 2,064 orthologous genes from the 13 schizothoracine fish. Of these, we identified 52 genes as candidate genes. These candidate genes have probably undergone positive selection along the whole schizothoracine fish lineage. We then found that nine candidate PSGs were significantly enriched in several key GO functions and metabolite pathways. Interestingly, all of the functions and metabolite pathways significantly enriched in the schizothoracine fish were associated with the immune system, which was inconsistent with previous studies of the schizothoracine fish (Yang et al. 2014; Tong et al. 2017). This discrepancy may be because previous studies focused on comparative transcriptomics between the schizothoracine fish and other teleost fishes from the plains, instead of interspecies comparisons within schizothoracine fish.

Fish immune function is known to be associated with habitat structure (Bowden 2008; Diepeveen et al. 2013). During the colonization of ecologically novel drainages or upon the exploitation of vacant ecological niches, selection on immune-related genes can be particularly strong as fishes encounter different pathogens (Birrer et al. 2012). Here, we identified nine candidate PSGs that were significantly enriched into five GO functions and two KEGG pathways. All of the enriched GO functions were related to the immune system, including cytokine activity, cytokine receptor binding, chemokine activity, chemokine receptor binding, and immune response. These results were supported by our KEGG pathway analysis.

Cytokines are a family of low-molecular-weight proteins that are secreted by activated immune-related cells upon induction by various parasitic, bacterial, or viral pathogens (Salazar-Mather and Hokeness 2006). Cytokines regulate the immune response as autocrines or paracrines by binding to the appropriate receptors (Zhu et al. 2013a). Cytokines are divided into interferons (IFNs), interleukins (ILs), tumor necrosis factors (TNFs), colony-stimulating factors, and chemokines; cytokines are secreted by various types of immune cells, including macrophages, lymphocytes, granulocytes, dendritic cells, mast cells, and epithelial cells (Salazar-Mather and Hokeness 2006). Therefore, cytokines have been catalogued as key regulators of the immune response, acting as a bridge between innate and adaptive responses (Alejo and Tafalla 2011). Here, seven of the nine candidate PSGs that were significantly enriched in GO functions and KEGG pathways were either cytokines or cytokine receptors, including interferon receptor 1 (IFNAR1, OG07729), C-C motif chemokine 25 (CCL25, OG11017), C-C motif chemokine 3 (CCL3, OG11002), C-C motif chemokine 8 (CCL8, OG03673), interleukin-34 (IL-34, OG11127), interleukin 2 receptor, gamma (IL2RG, OG09047), and interleukin-12 alpha chain (IL-12, OG06385).

Chemokines are early-response, chemotactic members of cytokine family secreted by infected tissue cells (Salazar-Mather and Hokeness 2006; Alejo and Tafalla 2011; Bird and Tafalla 2015). Chemokines recruit monocytes, neutrophils, and other effector cells from blood vessels to the infection site (Rajesh 2016). In fish, CCL3, CCL8, and CCL25 have been shown to play important roles in the recruitment of monocytes and leukocytes, as well as in the induction of the inflammatory response to microbial invasion (de Oliveira et al. 2013; de Oliveira et al. 2015).

Interleukins, the largest group of cytokines, are associated with both pro- and anti-inflammatory functions (Rajesh 2016). For example, IL-12 is produced primarily by antigen presenting cells (APC), such as macrophages, dendritic cells, and B cells; however, this cytokine is indispensable for macrophage, neutrophil, and lymphocyte recruitment to infected tissues and for the activation of these cells as pathogen eliminators (Svanborg et al. 1999). In fish, IL-34 expression is sensitive to inflammatory stimuli and may regulate macrophage biology (Wang et al. 2013).

Interferons (IFNs) are secreted proteins that may induce an antiviral state in cells, and that, in vertebrates, play a major role in antiviral defense (Samuel 2001). During viral infections, IFNs bind to the IFN receptors (IFNARs), which trigger signal transduction through the JAK-STAT signal transduction pathway, resulting in the expression of Mx and other antiviral proteins (Samuel 2001).

The schizothoracine fish are widely distributed in drainages throughout the QTP (Wu and Wu 1992; Chen and Cao 2000). Although there is little information about fish pathogens in the QTP, the different aquatic environments of the QTP may have different microbial populations and, thus, potentially unique pathogens. Even in the same drainage system, pathogens may be unique to a given micro-ecosystem due to differences in dissolved oxygen, ultraviolet radiation, and physical and chemical properties of the water. Thus, different schizothoracine fish are inevitably under selective pressures from new pathogens during the colonization of ecologically novel drainages or upon the exploitation of vacant ecological niches. Here, our lineage-specific test of the evolution of the nine candidate PSGs indicated that species from the same drainage shared more of the same PSGs. However, these PSGs also had species-specific features due to ecological differences among drainages, as well as among micro-habitats in the same drainage (e.g., benthic and pelagic). For example, of the four species collected from the Tsangpo River (*S. younghusbandi*, *P. dipogon*, *Schizothorax macropogon*, and *Oxygymnocypris stewartii*), three (*S. younghusbandi*, *P. dipogon*, and *S. macropogon*) are benthic (Wu and Wu 1992; Chen and Cao 2000; Qi et al. 2012). However, *S. younghusbandi* and *P. dipogon* share only eight PSGs, while *S. macropogon* shares only six. *O. stewartii*, a pelagic fish, shares four PSGs with *S. younghusbandi*, *P. dipogon*, and *S. macropogon*. Similar patterns were observed in the fish species from the Yellow River.

Fish are armed with less-developed adaptive immune systems as compared to those or mammals (Meng et al. 2012). The innate immune system plays an important role in fish, acting to rapidly eliminate pathogens, including bacteria and parasites, as the first line of defense against infection (Zhu et al. 2013b; Tong et al. 2017). The previous comparative transcriptomics of the schizothoracine fish and other teleost fish from the plains (Yang et al. 2014; Tong et al. 2017), together with our findings based on interspecific transcriptomic comparisons within the schizothoracine fish, suggested that genes associated with energy metabolism and the hypoxia response in the schizothoracine have been subjected to accelerated or adaptive evolution. These genes may have been critical for the adaptation of the schizothoracine fish to the cold and hypoxic environment of the uplifted QTP. In addition, the adaptive evolution of immune genes may have allowed the fish species to colonize ecologically novel environments or to exploit vacant ecological niches during speciation.

In summary, we assembled the transcriptomes of 13 schizothoracine fish. The genetic adaptation of these fish to local aquatic environments during speciation was investigated, based on an interspecific comparative analysis of the transcriptomes. We identified nine PSGs (out of 52) that were significantly enriched in immune system-related functions and metabolite pathways, which indicated that species-specific differences may have evolved in response to different ecological environments and to different living habits (e.g., benthic or pelagic). Our results suggested that the adaptive evolution of immune genes allowed new schizothoracine species to colonize ecologically novel environments or to exploit vacant ecological niches during speciation.

## ACKNOWLEDGMENTS

This work was supported by grants from the National Natural Science Foundation of China (31460094) and the Natural Science Foundation of Qinghai Science & Technology Department in China (2015-ZJ-901). We thank LetPub (www.letpub.com) for its linguistic assistance during the preparation of this manuscript.

